# Pupillometry and Whole-Brain c-Fos Mapping Uncover Multimodal Mirror Mechanisms in Emotional Contagion Networks of Mice

**DOI:** 10.1101/2024.11.20.624327

**Authors:** Matteo Caldarelli, Stefano Zucca, Aurelia Viglione, Alessandra Stella, Rida Nisar, Giulia Sagona, Ester Maria Papini, Fabio Carrara, Serena Bovetti, Raffaele Mazziotti, Tommaso Pizzorusso

**Author notes:** Corresponding author: Raffaele Mazziotti, Department of Neuroscience, Psychology, Pharmacology, and Child Health, University of Florence, Florence, ITALY. These authors contributed equally to this work (Matteo Caldarelli, Stefano Zucca, Aurelia Viglione).

## Abstract

Emotional contagion (ECo) represents a fundamental form of empathy. In this study, we used pupillometry to quantify ECo by assessing pupil responses of a mouse watching another mouse receive a tail shock. Pupil dilation effectively measured both direct and vicarious emotional response thresholds at the individual level through psychometric curve analysis. The pupillary ECo response diminished when the observer could not see the demonstrator, suggesting a multisensory process involving vision. Viewing videos of tail-shocked mice was sufficient to elicit a pupil response in the observer. Whole-brain c-Fos mapping revealed a broad network of 88 brain regions activated during ECo, with all areas activated in the demonstrator also engaged in the observer. Additionally, in certain brain regions, correlated activation was detected between each observer-demonstrator pair, indicating that ECo promotes a shared neural state. These findings advance our understanding of the neural basis of empathy, with implications for analyzing neuropsychiatric disorder models.

## Introduction

Empathy refers to the sharing of emotional states between individuals (Chang-Arana et al., 2022). This multifaceted concept spans from basic behavioral reactions to complex constructs like prosocial behavior, perspective-taking, and theory of mind (Cuff et al., 2014). Emotional contagion (ECo), one of the most simple forms of empathy, is an automatic emotional response to others emotions (Meyza & Knapska, 2018). Extensive research has been conducted on ECo in humans (Herrando & Constantinides, 2021; Meyza & Knapska, 2018) and animals (Pérez-Manrique & Gomila, 2022), revealing its presence in most of tested mammals and even in other groups such as in birds, like corvids (Adriaense et al., 2019; Wenig et al., 2021), fish (Burbano Lombana et al., 2021; Kareklas & Oliveira, 2024) and in much simpler animals such as isopods (Broly & Deneubourg, 2015). Several paradigms have been used to study ECo in rodents (Keysers et al., 2022; Kim et al., 2021; Yu et al., 2024), typically involving an observer and a demonstrator (Han et al., 2020). In the *Vicarious freezing* test, the observer increases freezing behavior in response to the negative emotions experienced by the demonstrator. In the *Vicarious Learning* paradigm, the observer learns new associations between stimuli by perceiving the emotional reactions of the demonstrator. These paradigms illustrate how ECo enables animals to avoid threats without direct exposure (Keysers et al., 2022).

ECo is an internal state (Keysers et al., 2022). Internal states involve dynamic variations in physiological and behavioral variables in response to stimuli. These pleiotropic effects consist of changes across different systems and occur in a coordinated manner. The extent of these fluctuations is proportional to the intensity of the stimulus (Flavell et al., 2022). Therefore, ECo is expected to share these characteristics, manifesting as widespread and concurrent changes across multiple physiological variables, with the magnitude of these responses corresponding to the emotional intensity evoked by the stimulus. The triggering of the pleiotropic effects involves modifications in the activity of the autonomic nervous system, which in turn are reflected on pupil size (Grujic et al., 2024; Viglione et al., 2023). For this reason pupillometry can be potentially considered as a window on the emotional system (Ferencová et al., 2021; Grujic et al., 2024; Kutlubaev et al., 2024; Nakakoga et al., 2020; Viglione et al., 2023). It has been recently shown that pupillary activity can address variations in the brain state not associated with noticeable changes in overt behavior (Ganea et al., 2020; Laeng et al., 2012; McGinley et al., 2015; Vinck et al., 2015). This indicates that pupillometry has high sensitivity to small and transient arousal fluctuations and it can detect both overt and covert behavioral phenomena. For these reasons, we evaluated the effectiveness of pupillometry in quantifying direct emotional response (DER) elicited by aversive stimulation, and vicarious emotional response (VER) elicited by witnessing the DER expressed by a conspecific.

Research has identified various brain areas involved in ECo in rodents, with most studies highlighting the significant role of the cortico-limbic circuit (Hernandez-Lallement et al., 2022; Keysers & Gazzola, 2023) and suggesting a significant overlap in neural activation between rodents and humans during ECo (Hernandez-Lallement et al., 2022). c-Fos activation was observed in several areas of the mouse brain such as the anterior cingulate cortex (ACA), prelimbic cortex (PL), and infralimbic cortex (IL), along with subcortical structures like the nucleus accumbens (NAc), mediodorsal thalamus (MD), lateral habenula (LH) and the amygdalar complex (Choi & Jeong, 2017; Kim et al., 2023; Mondoloni et al., 2024; Pisansky et al., 2017; Qi et al., 2022; Smith et al., 2021; Zheng et al., 2020). However, no studies have conducted a comprehensive analysis of whole-brain activation and network connectivity involved in ECo, nor have they explored how this activation relates to interactions between subjects in a dyad. In this work, we employed light-sheet microscopy to detect the areas activated during ECo, providing a detailed map of whole-brain activation. We then performed a connectivity analysis to study the underlying network of activated regions. Finally, we conducted a correlation analysis to determine if the activation patterns in the demonstrator are related to those in the observer, offering new insights into how activation may be coordinated between subjects in a dyad.

## Results

### Pupil size and locomotor activity are quantitative indicators of emotional reactivity to direct aversive stimulation

To assess whether pupillometry can detect shifts in internal states due to direct aversive stimulation, we developed a paradigm where head-fixed mice are subjected to tail shocks of varying intensities while simultaneously monitoring both pupillary and locomotor activity (Fig. 1a). Tail shocks resulted in a significant transient increase in both pupil dilation and locomotor activity across all intensities relative to baseline (Fig. 1b-c). Moreover, the magnitude of these responses scaled proportionally with the intensity of the stimulation (Fig. 1d). Under our experimental conditions, mice responded to tail shocks by fleeing, while other studies report freezing behavior in response to aversive stimuli. This variation aligns with previous literature, which shows that whether a mouse flees or freezes in response to a threatening stimulus depends on environmental, situational, and biological factors (De Franceschi et al., 2016; Eilam, 2005; Hébert et al., 2019). These findings indicate that pupillometry, alongside locomotor activity, provides a reliable marker of direct emotional responses to aversive stimuli, with both pupil dilation and locomotion reflecting the subjective experience of the stimulus.

**Figure 1.**
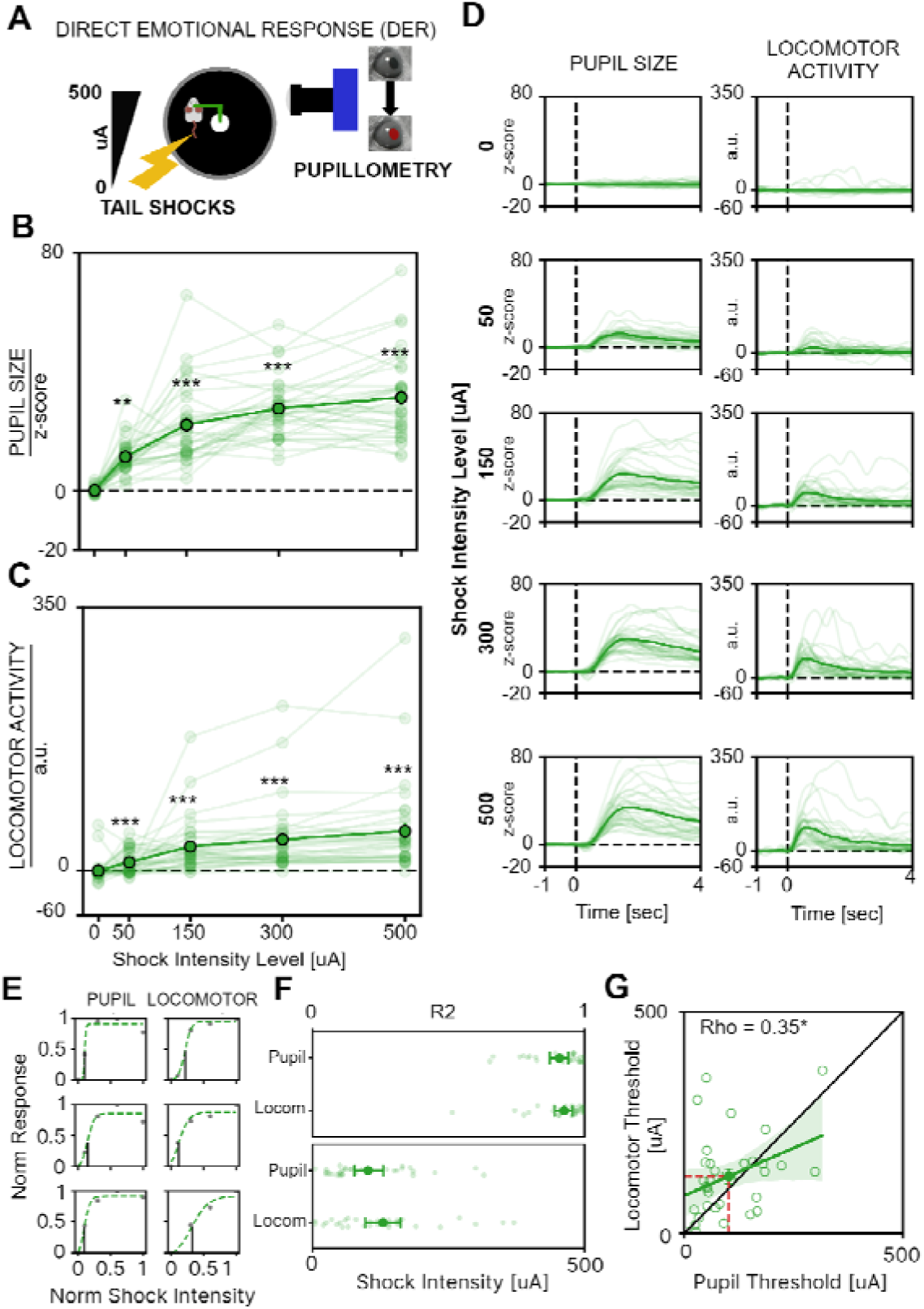
Effects of DER on Pupillary and Locomotor Activity in Mice. **A)** Sketch of the experimental setup: head-fixed mice on a circular treadmill receive tail shocks ranging from 0 to 500 µA. **B)** Average pupillary dilation as a function of shock amplitude. Thin lines represent individual mouse traces, thick lines indicate the average response acros all animals (rm one-way ANOVA, n=32, F_4,124_=17.89, p<0.001, followed by BH procedure for multiple comparisons; asterisks represent significance vs 0 µA intensity stimulation). **C)** Average locomotor activity as a function of shock amplitude. Thin lines represent individual mouse traces, thick lines indicate the average response acros all animals (rm one-way ANOVA, n=32, F_4,124_=76.36, p<0.001, followed by BH procedure for multiple comparisons; asterisks represent significance vs 0 µA intensity stimulation). **D)** Grand average of the temporal profile of pupil size (left) and locomotor activity (right) at different stimulation intensities. The vertical dashed line marks the onset of the stimulus. Thin lines depict average responses of each subject. **E)** Representative single-animal pupillary and locomotor responses fitted by psychometric curves. Solid gray circles represent raw responses, and the dashed green lines indicate the fitted curves. Vertical black lines mark the threshold level for each psychometric curve, representing the intensity value at 50% of the maximal response. **F)** Top: goodness of psychometric curve fitting (R^2^) of the data of each mouse. Bottom: response thresholds for every mouse. Light points represent single animal thresholds, solid points represent mean ± SEM. **G)** Correlation between locomotor and pupil thresholds. Thick line indicates correlation line, shaded area is 95% CI. Data are presented as mean ± SEM (Spearman’s correlation, n=32, ρ=0.35, p<0.05). (*: p<0.05. **: p<0.01, ***: p<0.001).

To further investigate the capacity of pupillary and locomotor activity to profile individual sensitivity to aversive stimulation, we applied psychometric fitting to the responses of individual animals (Fig. 1e). This method allows us to quantify how each DER of the subject scales with stimulus intensity, providing parameters that describe the performance. The goodness-of-fit metric demonstrates that psychometric fitting accurately models both pupillary and locomotor response curves, with no significant differences in fit quality or average responsiveness thresholds between the two measures (Fig. 1f). No difference in DER threshold was present between male and female mice (Fig. S1d). We also found that pupillary and locomotor response thresholds are correlated within subjects, suggesting that both pupillary and locomotor activity can effectively quantify the emotional reactivity of the subject to aversive events (Fig. 1g).

### Pupil size, but not locomotor activity, is a quantitative indicator of rapid vicarious emotional response

To evaluate whether pupillary and locomotor activity can track ECo in the observing mouse, we assessed VER using two mice: an observer and a demonstrator. Both mice were head-fixed and maintained in multisensory contact, allowing for auditory, visual, and olfactory interaction (Fig. 2a). The demonstrator received tail shocks of varying intensities as shown in Fig. 2a, while the pupillary and locomotor activity of the observer were recorded. We found that when the demonstrator received a tail shock, the observer exhibited a significant pupil dilation, while no changes in locomotor responses were detected (Fig. 2b-d). These findings demonstrate that pupil size measures can effectively track VER occurring in the observer mouse. Furthermore, applying psychometric fitting to pupillary VER of individual observers revealed a good fit quality, comparable to that observed measuring DER (Fig. 2e-f). This suggests that psychometric fitting of pupillary VER is a reliable method for assessing ECo thresholds at the single-subject level. No difference in VER threshold was present between male and female mice (Fig. S1d). Notably, the VER threshold was significantly higher than the DER threshold (Fig. 2f). The analysis of VER and DER thresholds in the same animal did not show a significant correlation between these measures, suggesting that the processes determining the sensitivity to direct and vicarious emotional response can be dissociated (Fig. 2g).

**Figure 2.**
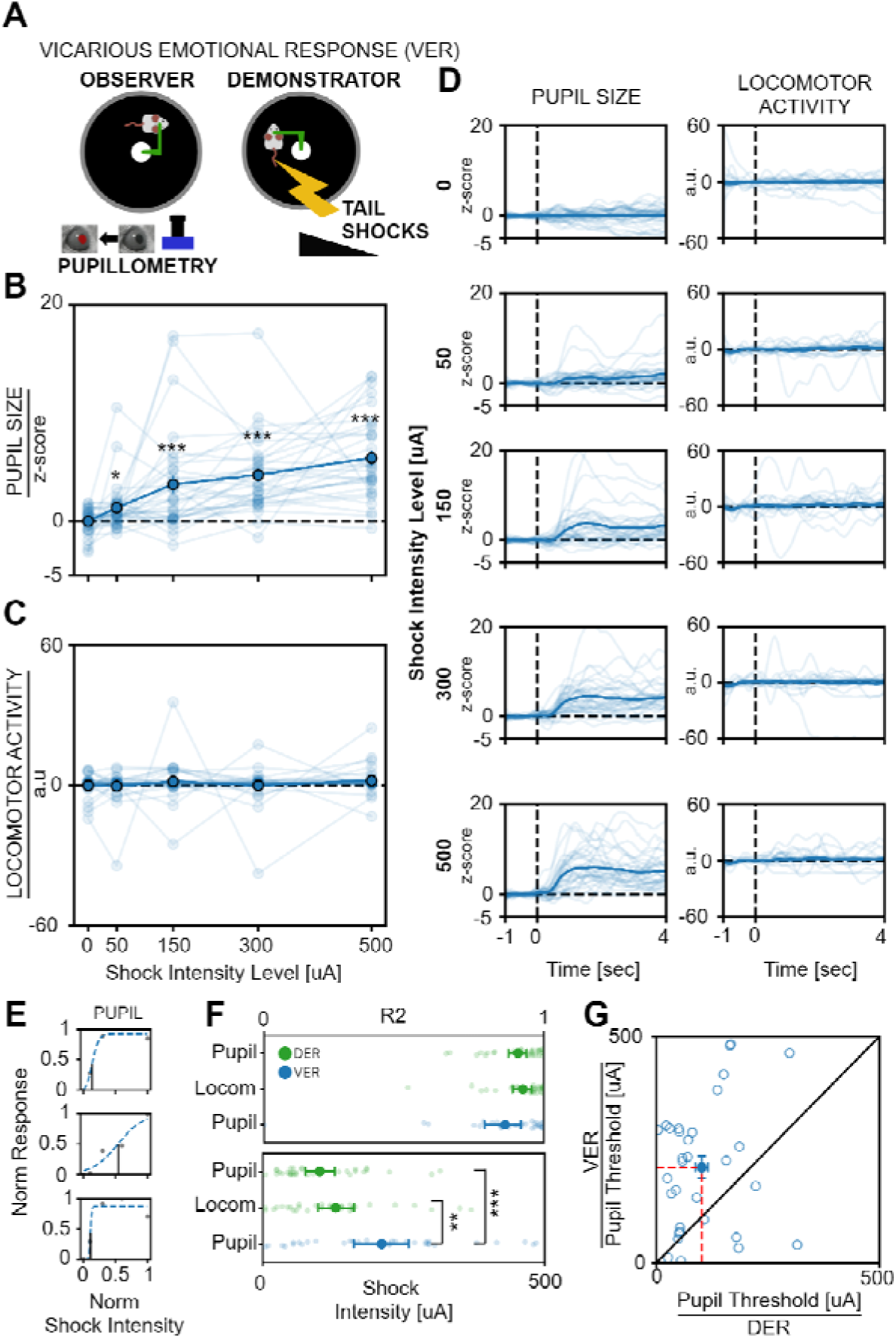
Effects of VER on Pupillary and Locomotor Activity in Mice. **A)** Sketch of the experimental setup: two head-fixed mice are placed on separate circular treadmills, allowing visual, olfactory, and auditory sensory interaction. Tail shocks ranging from 0 to 500 µA are applied to the demonstrator, while recording pupillary responses and locomotor activity in the observer. **B)** Average pupillary dilation of the observer as a function of the shock amplitude applied to the demonstrator. Thin lines represent individual mouse traces, thick lines indicate the average response across all animals (rm one-way ANOVA, n=32, F_4,124_=20.27, p<0.001 followed by BH procedure for multiple comparisons; asterisk represent significance vs 0 µA intensity stimulation). **C)** Average locomotor activity of the observer as a function of the shock amplitude applied to the demonstrator. Thin lines represent individual observer responses, the thick line represents the average response (rm one-way ANOVA, n=32, F_4,124_=0.83, p=0.51, followed by BH procedure for multiple comparisons). **D)** Grand average of the temporal profile of pupil size (left) and locomotor activity (right) at different stimulation intensities. The vertical dashed line marks the onset of the stimulus. Thin lines depict average responses of each subject. **E)** Representative single-animal pupillary and locomotor responses fitted by psychometric curves for VER. Solid gray circles represent raw responses, and the dashed green lines indicate the fitted curves. Vertical black lines mark the threshold level for each psychometric curve, representing the intensity value at 50% of the maximal response. **F)** Top: Goodness of psychometric curve fitting for VER (pupil) and DER (pupil and locomotor activity) does not differ (one-way ANOVA, n=32, F_2, 93_=1.83, p=0.17). Bottom: Comparison of response thresholds between VER (pupil) and DER (pupil and locomotor activity) conditions (one-way ANOVA, n=32, F_2, 93_=8.95, p<0.01, followed by BH procedure for multiple comparisons). **G)** No correlation between pupil response thresholds of observers is present during a VER or a DER. Data are presented as mean ± SEM (Spearman’s correlation, n=32, ρ=0.075, p=0.68). (*: p<0.05. **: p<0.01, ***: p<0.001).

### Visual input contributes to ECo

Previous research indicates that ECo is a multimodal process, with most studies focusing on the contribution of olfactory and auditory stimuli (Hernandez-Lallement et al., 2022), however little is known about the role of vision in ECo. To address this question, we implemented two modified versions of the VER experiment: the VER+occluder condition, in which an opaque screen was positioned between the animals to obstruct the observer’s view, aimed at determining whether ECo operates via non-visual sensory modalities; and the Vision-only condition, where the demonstrator was replaced by a video of a mouse undergoing direct aversive stimulation, to assess the contribution of visual cues to ECo (Fig. 3a). The results were compared to those of mice performing the standard VER experiment, as depicted in Fig. 2.

**Figure 3.**
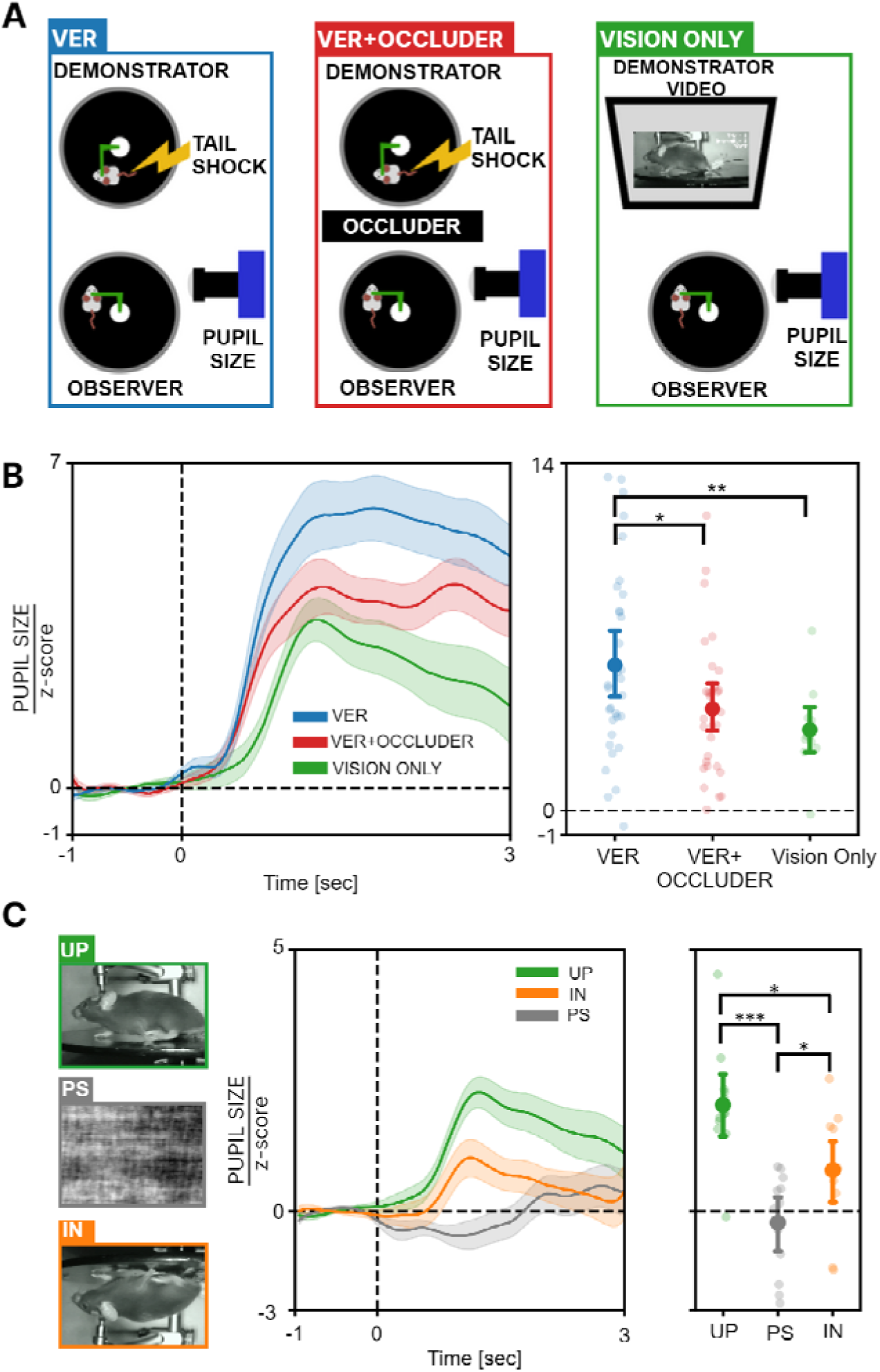
Activation of Pupil VER is a Multisensory Process that also Involves Vision. **A)** Sketch of the experimental setting. In the VER condition, the observer and demonstrator are in multisensory contact. In the VER+Occluder condition, a barrier is placed between the observer and demonstrator and shocks of 0 or 500 uA are delivered to the demonstrator. In the Vision Only condition, the observer mouse watches a video of a demonstrator receiving either 0 or 500 µA shocks. **B)** Left: average time course of the pupillary responses in the VER, VER+Occluder and Vision Onl conditions. Right: average pupillary dilation in VER, VER+Occluder and Vision Onl conditions. Solid circles represent the average and lighter circles represent single animal data. Data are presented as the mean ± 95% confidence interval (one-wa ANOVA, n_VER_=32, n_VER+OCCLUDER_=30, n_VISIO ONLY_=12, F_2, 71_=3.91, p<0.05, followed by BH procedure for multiple comparisons). **C)** Left: example of frames from movies used to induce ECo (UP: video of a mouse receiving the shock, PS: phase scrambled version of the same movie, IN: inverted version of the same movie). Center: time course of pupil size across the three conditions. Right: pupillary dilation in UP, IN and PS conditions. Solid points represent the average and lighter points represent single animal data. Data are presented as the mean ± 95% confidence interval (one-way ANOVA, n=12, F_2, 22_=18.42, p<0.001, followed by BH procedure for multiple comparisons). (*: p<0.05. **: p<0.01, ***: p<0.001).

We found that the VER+occluder condition evoked a smaller, but significant pupil dilation as compared to the response to the VER condition (Fig. 3b). This indicates that although vision is not strictly required to elicit a vicarious pupillary response, it significantly modulates its amplitude. This observation suggests that ECo processing undergoes multisensory enhancement, wherein the integration of multiple sensory inputs leads to a more pronounced response, amplifying the pupillary event (Klasen et al., 2012).

Strikingly, also the Vision Only condition elicited a significant pupil dilation demonstrating that the visual modality alone is sufficient to reliably trigger a VER, albeit of reduced amplitude with respect to a multisensory VER. To further explore the nature of the pupillary response in Vision Only mice, we introduced additional variations to the protocol aiming at dissecting the specific contributions of visual processing. Specifically, we generated two versions of the original Upright (UP) video to assess the effect of two distinct visual stimulations. The first variation, termed Phase-scrambled (PS), involved scrambling the Fourier phase of the video frames, preserving key physical parameters such as luminance, contrast, and motion, while disrupting the semantic integrity of the images. The second variation, referred to as Inverted (IN), employed a vertical flip of the video frames, maintaining the physical properties but altering the spatial configuration. Our results revealed that the IN condition elicited a pupillary response exceeding the PS condition, although this response was significantly attenuated compared to the UP condition (Fig. 3c). These findings suggest that observer mice exhibit a preference for social stimuli with the expected spatial configuration, implying that the pupillary response is not driven solely by alterations in physical image properties. This indicates the presence of holistic processing of mouse body configurations, similar to the mechanisms underlying facial emotional recognition in humans (Pallett & Meng, 2015; Poltoratski et al., 2021).

### Whole brain c-Fos analysis reveals a large, partially overlapping, and distributed network of activation in response to direct and vicarious emotional stimulation

To clarify the neural circuits activated during DER and VER, we conducted whole-brain analysis of c-Fos activation in the observer and the demonstrator with respect to control mice. Mice belonging to the three groups were sacrificed 90 minutes after the end of stimulation and c-Fos activation was assessed by whole-brain immunolabeling, iDISCO tissue clearing and light-sheet microscopy (Fig. 4a-b). The control mice were head-fixed in the same setup for the same duration of experimental mice but did not receive shocks or observe any shocked mouse. ClearMap2 toolbox (Kirst et al., 2020) was used to align the images to the Allen Atlas Mouse Brain (25µm -v2) and to perform automatic whole-brain detection of c-Fos^+^ cells in 169 regions across the entire brain (see materials and methods for details). To avoid errors due to mis-alignment or tissue damage, the medulla, the pons and the cerebellum were excluded from the analysis.

**Figure 4.**
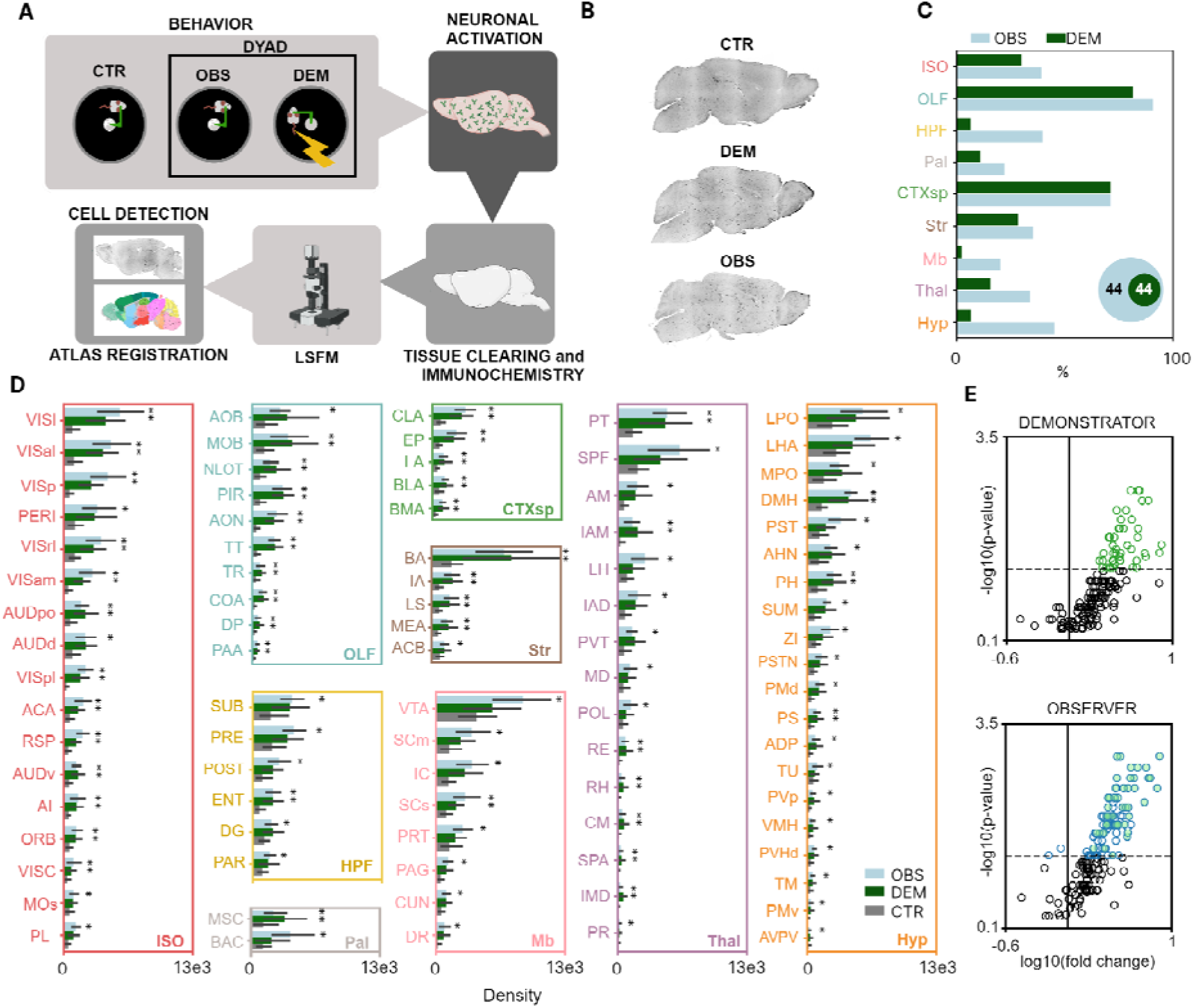
Whole-brain c-Fos Mapping. **A)** Experimental pipeline for whole-brain c-Fos mapping of recruited brain areas: animals from the three different groups (control, CTR; demonstrator, DEM; and observer, OBS). **B)** Representative images of c-Fos stained tissues in the three different conditions. **C)** Percentage of brain areas with significant c-fos activation in the OBS (light blue) and DEM (green) group compared to the CTR (gray) group, relative to the total number of regions defined in each Allen Brain Atlas macroarea. (ISO: isocortex, OLF: olfactory areas, HPF: hippocampal formation, CTXsp: cortical subplate, Str: striatum, Pal: pallidum, Thal: thalamus, Hyp: hypothalamus, and Mb: midbrain). <u>Inset</u>: Venn Diagrams illustrating the number of significantly activated brain areas in OBS and DEM groups compared to CTR. **D)** Bar graphs showing c-Fos^+^ cell density across the three experimental groups in brain regions with significant modulation in either the OBS or DEM conditions compared to the CTR group. Values are presented as the mean ± 95% confidence interval, with brain regions grouped according to their macro-area classification in the Allen Brain Atlas hierarchy (Mann-Whitney U Test, *: p<0.05; n_OBS_ = 7, n_DEM_ = 7, n_CTR_ = 6). **E)** Volcano plots of the c-Fos^+^ cell density fold change in the DEM (upper) or OBS (lower) condition calculated over the CTR group across all brain regions (empty circles). Colored circles indicate significantly modulated areas. Acronyms for brain areas are defined in Supplementary Table 2.

We identified 88 brain regions significantly modulated in either the observer or the demonstrator group compared to the control group (Fig. 4c-e, Fig. S2c and Supplementary Table T1). These regions spanned nearly all major brain structures, with a higher proportion of recruited areas in the observer group compared to the demonstrator across all primary structures except the cortical subplate (Fig. 4c). Interestingly, 44 out of 88 regions showed c-Fos upregulation exclusively in the observer group, while every region activated in the demonstrator group was also recruited in the observer (Fig. 4c-d). This pattern suggests the presence of two co-activated networks: one specific to ECo, and another shared circuit engaged in processing aversive emotional stimuli. Areas previously known to be involved in ECo (Hernandez-Lallement et al., 2022; Keysers et al., 2022; Smith et al., 2021) such as the anterior cingulate cortex (ACA), agranular insular area (AI), nucleus accumbens (ACB), mediodorsal thalamus (MD), lateral habenula (LH), medial amygdala (MEA), basolateral amygdala (BLA), zona incerta (ZI), ventral tegmental area (VTA) and periaqueductal gray (PAG) were significantly activated in our analysis.

Since values of c-Fos^+^ cell density provide a single activation index for each region per animal, we then performed an estimation of the functional connectivity by calculating the covariance across subjects belonging to the same experimental group and across all pairs of brain areas. We found an overall stronger correlation in the demonstrator group compared to the observer and control conditions (Fig. S2a; median value: 0.75 for demonstrator, 0.4176 for observer and 0.4385 in control, p<0.001, Kruskal-Wallis Test), indicating that, despite the recruitment of partially overlapped brain circuits, the overall network is functionally different under the two conditions.

To investigate the anatomical connectivity patterns between the recruited brain regions, we constructed an anatomical connectome using data from the Allen Brain Connectivity Atlas (Knox et al., 2019) generating a weighted, directed matrix representing connection strengths between the activated regions (Fig. 5a). We then applied the Leiden algorithm (Traag et al., 2019) to partition the network based on connection density. This analysis identified five clusters of interconnected regions, highlighting the organization of the network based on connection density (Fig. 5b) and the complex multisensory bases of ECo. A first cluster was purely hippocampal, a second cluster included mostly thalamic and prefrontal cortical areas; a third cluster included sensory subcortical and cortical areas; a fourth cluster included areas important for emotional control such as parts of the amygdala, the insular and visceral cortex, and olfactory areas; and a fifth cluster containing mainly hypothalamic areas activated only in the observer group.

**Figure 5.**
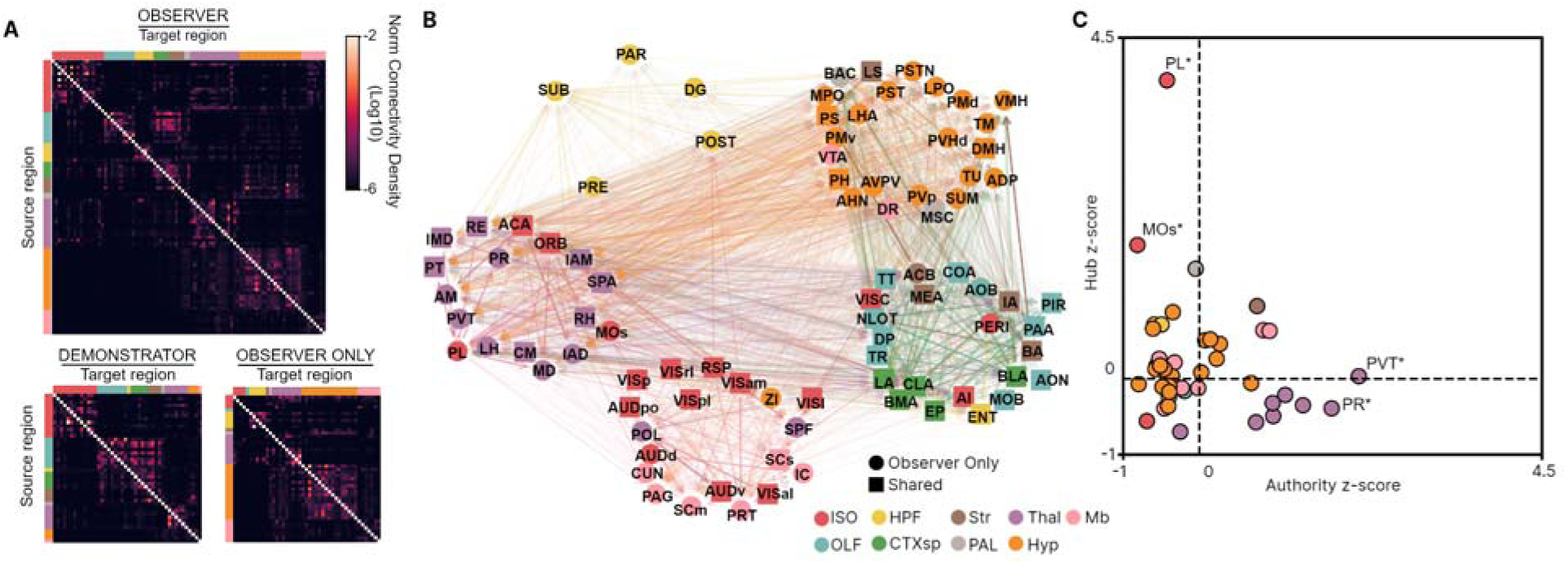
Network Analysis of Activated Areas. **A)** Weighted-directed matrices of anatomical connectivity between brain areas significantly activated in observers (top), in demonstrators (bottom-left), and exclusively in observers (bottom-right) **B)** Partitioned network of activated areas based on the anatomical connectivity. Line opacity reflects the strength of connections, while arrows indicate directionality. Circles represent areas activated only in observers, and squares represent areas shared between observers and demonstrators. **C)** HITS analysis of areas activated exclusively in observer (Bootstrap procedure, n_iterations_=1000, *: p<0.05 show a statistically significant Hub-score for PL and MOs, or a significant Authority-score for PR and PVT). Areas are color-coded according to the macro-region they belong to; ISO: isocortex, OLF: olfactory areas, HPF: hippocampal formation, CTXsp: cortical subplate, Str: striatum, Pal: pallidum, Thal: thalamus, Hyp: hypothalamus, and Mb: midbrain. Acronym for brain areas are defined in Supplementary Table 2.

Given that the observer group exhibits a unique array of brain regions activated specifically during ECo, we conducted a more detailed analysis of the connectivity patterns within this circuit to better understand the role of each region in the network. Specifically, we quantified the roles of individual regions as hubs and authorities within the network using the Hyperlink-Induced Topic Search (HITS) algorithm (Kleinberg, 1999). An authority is defined as a node that integrates a significant amount of information from other key nodes, while a hub is a node that transmits information to other nodes of the network. To calculate the significance level of each area as a hub or authority in the observed network, we compared the values to the average hub and authority scores obtained using random networks generated via bootstrapping (see Methods). Fig. 5c shows that the prelimbic cortex (PL) and secondary motor area (MOs) emerge as key hubs, while the paraventricular nucleus of the thalamus (PVT) and perireunensis nucleus (PR) are key authorities.

### Coupled pupillary and neural responses in dyads of observer and demonstrator mice

ECo is a process observed in individuals exposed to the emotions of others, leading members of a dyad to potentially exhibit similar behavioral patterns. To test this possibility, we first analyzed the correlation between the pupillary responses of the observer and demonstrator within each dyad (Fig. 6a). Our findings revealed a positive correlation between the pupil dilation thresholds of observers and demonstrators within each dyad (Fig. 6b). As expected, this correlation was not detected when comparing pupil dilation thresholds of observers with locomotor activity thresholds of demonstrators (Fig. 6c). This result suggests that the threshold for ECo of the observer mouse is a function of the internal state of the demonstrator. To study in depth this possibility we asked if the link between subjects of the same dyad could also be reflected in the activation of brain areas. We thus assessed the correlation of c-Fos^+^ cell density for all dyads restricting the analysis to the 44 brain areas which were significantly recruited in both the observer and demonstrator groups. We compared the results with a surrogate dataset generated by shuffling the animal pairs across dyads and we only considered areas showing a correlation coefficient higher than the 95th percentile of the shuffled distribution. Strikingly, a similar observer-demonstrator dyad correlation was identified in the c-Fos activation. In particular, the strongest association was observed in 10 brain areas, including the MEA, BLA, the sensory related superior colliculus (SCs), and AI (Fig. 6d). These regions are critical for processing emotional stimuli and regulating arousal (Gogolla, 2017; Smith et al., 2021; Solié et al., 2022), further supporting the link between shared emotional experiences and neural activation in the dyad.

**Figure 6.**
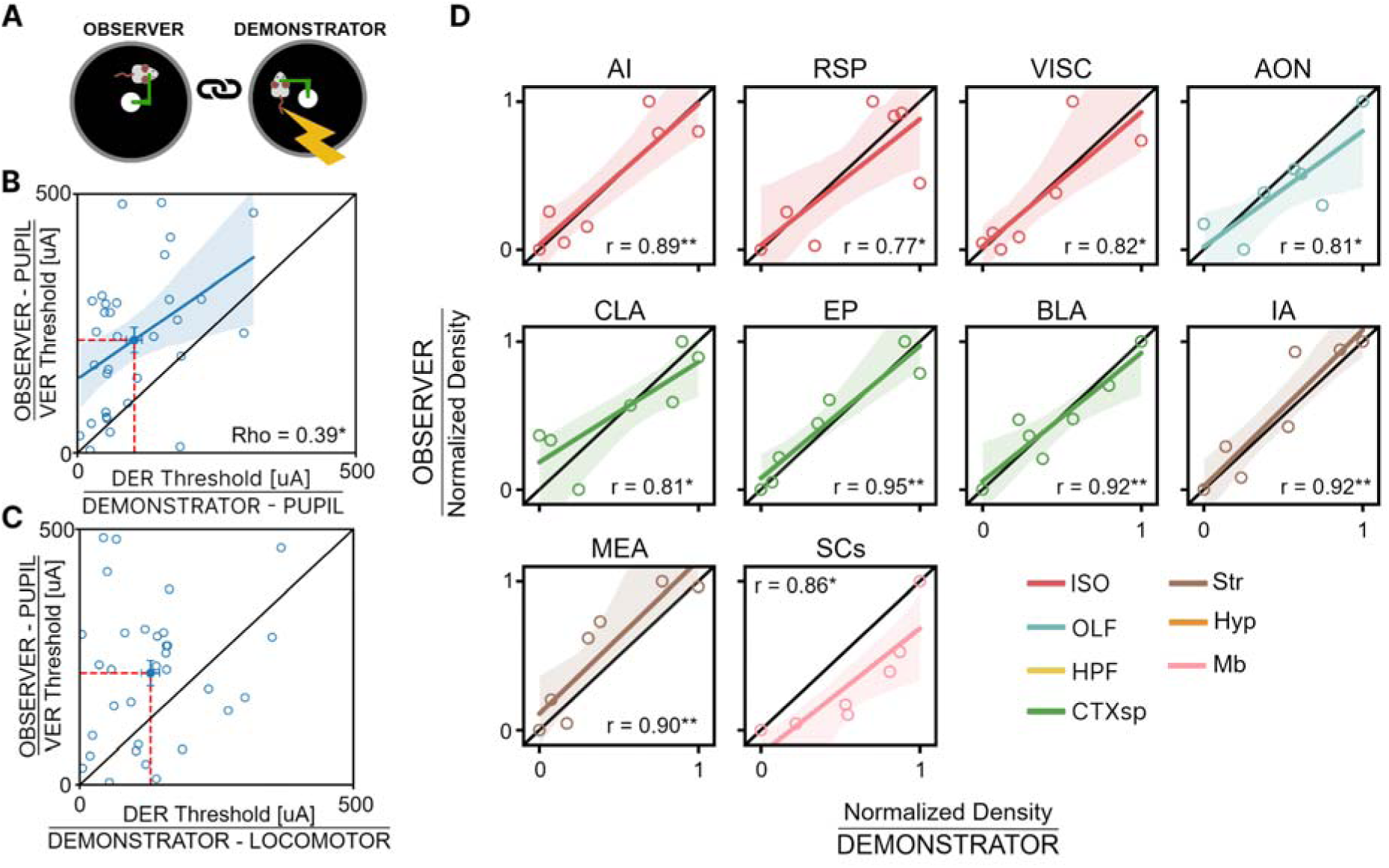
Within Dyad Correlation in Emotional Responses and c-fos activation. **A)** Sketch of a dyad. **B)** Correlation between the pupil VER thresholds of observers and the pupil DER thresholds of demonstrators in dyads. Data are presented as mean ± CI (Spearman’s correlation ± confidence interval, n=32, ρ=0.39, p<0.05). **C)** Correlation between the pupil VER thresholds of observers and the locomotor DER thresholds of demonstrators in dyads. Data are presented as mean ± SEM (Spearman’s correlation, n=32, ρ=0.13, p=0.46). **D)** Correlation between the c-fos activation of observers and demonstrators in dyads. Only areas showing significant correlation are reported. Data are presented as mean ± CI (Pearson’s correlation, n=7, correlation coefficients shown in figure, see Supplementary table T2 for areas nomenclature; areas are color-coded according to the macro-region they belong to; ISO: isocortex, OLF: olfactory areas, HPF: hippocampal formation, CTXsp: cortical subplate, Str: striatum, Pal: pallidum, Thal: thalamus, Hyp: hypothalamus, and Mb: midbrain). (*: p<0.05. **: p<0.01).

## Discussion

Our results demonstrate that pupil dynamics are a reliable and sensitive indicator of both direct and vicarious emotional reactivity to aversive stimulation. However, while both pupil dilation and locomotor activity were correlated with the intensity of the response to direct aversive stimulation, only pupillary responses were selectively triggered during ECo. Thus, pupil dilation during ECo can reveal forms of covert responses that cannot be detected with behavioral analysis. Importantly, pupillary DER and VER were accurately fitted by psychometric functions determining ECo sensitivity at single subject level. Altogether, these results highlight the usefulness of pupillometry as a non-invasive, highly translational and objective quantification of emotional responses.

### Neural Mechanisms of Direct and Vicarious Emotional Responses

Whole-brain c-Fos analysis revealed that while 44 regions were activated in demonstrators during aversive stimulation, a broader network of 88 regions was activated in the observers during ECo. While some of these areas were previously described to be involved in ECo (Hernandez-Lallement et al., 2022), our analysis indicates that a relatively large network is activated in the observer brain. Importantly, the observer network includes all of the regions activated in the demonstrator, indicating an overlap between the circuits engaged in both aversive stimulation and ECo. These findings raise interesting parallels with the mirror neuron system, which has been extensively studied for its role in action recognition and imitation (Di Cesare et al., 2015; Rizzolatti et al., 1996). While traditionally associated with motor functions, the mirror system has also been implicated in emotional recognition and empathy (Bekkali et al., 2021; Plata-Bello et al., 2023; Wu et al., 2023). The overlap between the regions activated during direct aversive stimulation in the demonstrator and ECo in the observer suggests that ECo could involve a similar mirroring process. This mirroring could extend beyond the motor system to facilitate emotional and affective processes potentially mediated by subcortical regions such as the amygdala and hypothalamus, which are key nodes in the regulation of emotional states.

The strong coupling between the brain activation of the observer and the demonstrator is also prompted by the strong correlation between c-Fos activation in some regions of the observer and demonstrator animals belonging to the same dyad. Notably, 10 specific brain regions, including the medial and basolateral amygdala, the superior colliculus, and insular cortex, showed significant correlations in activation levels between the subjects within the dyad. These regions are known to be involved in processing social and emotional stimuli, and regulating arousal (Klein et al., 2021; Nicolas et al., 2023; Solié et al., 2022), suggesting that the emotional states of the demonstrator and the observer are closely aligned at neural level. This activation coupling between demonstrator and observer may underlie the synchronization of emotional responses seen in ECo, further supporting the idea that emotional contagion involves a shared neural substrate between individuals (Prochazkova & Kret, 2017; Wang et al., 2024).

Our analysis demonstrated that both subcortical and cortical regions associated with visual, sensory, and olfactory systems are activated by ECo. This aligns with the hypothesis that ECo is driven by multisensory integration of various sensory inputs (Hernandez-Lallement et al., 2022). Our pupil analysis also supports this view showing that a VER can also be elicited in the absence of visual contact between the observer and the demonstrator, possibly by olfactory and auditory cues. However, the pupillary VER induced in this condition was smaller than normal, and videos representing tail-shocked demonstrators could evoke a pupillary VER. The visually induced VER was smaller than normal indicating that multisensory integration is needed to achieve a full scale VER. Notably, the detection of the demonstrator’s aversive response was highly specific to the spatial configuration of the image and could not be replicated by videos with similar visual characteristics but altered spatial configuration. Although it is well established that mice can discriminate conspecifics based on their emotional state (Jabarin et al., 2021; Scheggia et al., 2019), our findings suggest that this ability may involve a specialized system for visually recognizing emotions in conspecifics.

VER elicited a broader activation of brain regions in the observer compared to the demonstrator, which may subserve the processing of social and emotional signals. This suggests that the observer’s brain not only mirrors the emotional state of the demonstrator but also engages in higher-order processes linked to ECo (Lamm et al., 2016). The activation of this extended neural network in the observer could be in line with psychometric analyses of pupillary responses, which revealed significantly higher thresholds for VER compared to DER. Together, these findings offer a potential neural mechanism for understanding how emotional contagion contributes to empathy and social bonding in both animals and humans.

### Emotional Contagion in Neurodevelopmental and Neurodegenerative Conditions: Implications for Social and Cognitive Disorders

Our findings have significant implications for understanding social and emotional deficits in neurodevelopmental disorders such as autism spectrum disorder (ASD) and neurodegenerative conditions like Alzheimer’s disease (AD). ECo is a fundamental component of social interaction and empathy, both of which are dysregulated in ASD, where ECo is often decreased, leading to diminished emotional resonance (Harmsen, 2019; Helt et al., 2020; Kimmig et al., 2024). In contrast, emotional contagion is heightened in individuals with AD, possibly as a compensatory mechanism in response to cognitive decline, and it may serve as an early biomarker of the disease (Bible, 2013; Choi & Jeong, 2017; Giacomucci et al., 2024; Warren, 2022). The broader network activation observed in ECo suggests that these processes go beyond basic emotional responses. Deficits or alterations in this network may contribute to the difficulties in social communication and emotional resonance seen in ASD and AD, respectively. The identification of brain regions involved in emotional processing and ECo offers promising targets for studying how genetic mutations or neurodegenerative changes affect these circuits, which could guide interventions. Additionally, the use of pupillometry as a non-invasive marker of emotional reactivity provides opportunities for detecting subtle changes in emotional processing in patients and animal models of both ASD and AD. In conclusion, our findings highlight that pupillary responses offer a reliable window into the shared emotional states of empathy, revealing a multisensory process at work, where emotional contagion not only reaches across brains, but unites them in synchronized neural activity.

## Materials and Methods

### Animal handling

Animals were maintained in rooms at 22°C with a standard 12-h light-dark cycle. During the light phase, a constant illumination below 40 lux from fluorescent lamps was maintained. Food (standard diet, 4RF25 GLP Certificate, Mucedola) and water were available ad libitum and changed weekly. Open-top cages with wooden dust-free bedding were used. All the experiments were carried out according to the directives of the European Community Council (2011/63/EU) and approved by the Italian Ministry of Health (aut n. 357/2024-PR; prot. B4BB8.51). All necessary efforts were made to minimize both stress and the number of animals used. Weaning was performed on postnatal day (P)21–23. We tested male and female wild-type C57BL/6J (from P60 to P120; Charles River).

### Surgery

Mice were deeply anesthetized using isoflurane (3% induction and 1.5% maintenance), placed on a stereotaxic frame and head-fixed using ear bars. Prilocaine was used as a local anesthetic for the acoustic meatus. Body temperature was maintained at 37° C using a heating pad. The eyes were treated with a dexamethasone-based ophthalmic ointment (Tobradex, Alcon Novartis) to prevent cataract formation and keep the cornea moist. Respiration rate and pedal reflex were checked periodically to maintain an optimal level of anesthesia. Prior to scalp removal, a subcutaneous local injection of lidocaine (2%) was performed. After scalp removal, the skull surface was carefully cleaned with saline solution. After it dried, a thin layer of cyanoacrylate was poured over the exposed bone to attach a custom-made head post that was composed of a 3D printed base equipped with a glued set screw (12 mm long, M4 thread, Thorlabs: SS4MS12). The implant was secured to the skull using cyanoacrylate and ultraviolet curing dental cement (Fill Dent, Bludental). At the end of the surgical procedure, mice were left in a heated cage for recovery. After 1 h, mice were returned to their home cage. As antalgic therapy, paracetamol (100 mg/kg) was dissolved in drinking water for 3 days. At least 7 days were provided for the animals to fully recover before beginning of habituation procedures.

### Pupillometry and locomotor activity recordings

For pupillary and locomotor activity recordings, mice were head-fixed and free to run on a circular treadmill. A modified version of the apparatus proposed by Silasi et al. (Silasi et al., 2016) equipped with a 3D printed circular treadmill (⌀ 18 cm) was employed. Locomotor activity was detected using a rotary encoder (E38S6G5-600B-G24N). A USB camera (oCam-5CRO-U, Withrobot Lens: M12 25 mm) connected to a Jetson AGX Xavier Developer Kit (NVIDIA) running a custom Python3 script (30fps) was used to record the pupil. Real-time pupillometry was performed using MEYE (Mazziotti et al., 2021). To ensure a uniform light on the pupil an IR illuminator was used. Before the experiments, mice were handled for 1 week for 5 minutes each day; after, they were gradually introduced to head-fixation for an increasing amount of time for at least 5 days (habituation). During days 1 and 2, at least two sessions of 10 minutes of head-fixation, one in the morning and one in the afternoon, were performed. On Day 3, two sessions of 20 minutes. On day 4 one session of 40 minutes. On day 5 the last session was conducted for a time of 60 minutes. During each head-fixation session, a curved monitor (24 inches Samsung, CF390) was placed in front of the animal (at a distance of 13 cm), showing a uniform gray with a mean luminance of 8.5 cd/m^2^. A stimulation electrode, not connected to the stimulus isolator, was attached to the base of the mouse tail.

### Behavioral assessments

Aversive stimulation assessment and video administration experiments were performed in the same context and setup used for head-fixation habituation period. Emotional contagion assessment was performed in the same head-fixation setup but in a different context.

### Direct emotional response assessment

To assess pupillary and locomotor responses to the delivery of tail shocks, head-fixed mice were placed at a distance of 13 cm from the curved monitor showing a uniform gray with a mean luminance of 8.5 cd/m^2^. A stimulation electrode was attached to the base of the mouse tail and plugged to a stimulus isolator (World Precision Instruments, A320R). Mice were able to run on the circular treadmill. Pupillary area and locomotor activity were continuously recorded during the experiment. An ascending and then descending ladder of electric stimuli (0 µA, 50 µA, 150 µA, 300 µA, 500 µA, 0 µA, 500 µA, 300 µA, 150 µA, 50 µA; duration of 0.2 sec, interstimulus time of 60 sec) was delivered to mice. The ascending-descending ladder was repeated 3 times with a total of 6 repetitions for each stimulation level. The total duration of the experiment was 30-35 min.

### Vicarious emotional response assessment

In this experiment, littermate mice were used. One mouse was designated as the observer and the other as the demonstrator. The experiment was performed 48 hours after the aversive stimulation assessment. After another 48 hours, the experiment was repeated with the same pair of animals, with their roles reversed. Both mice ran on the circular treadmill. A stimulation electrode was attached to the tail of both animals, but only the electrode of the demonstrator was connected to the stimulus isolator. The pupillary size of the observer and the locomotor activity of both mice were continuously recorded during the experiment.

The demonstrator mouse was placed at the same height but perpendicular to the line of sight of the observer mouse, with its face centered in the binocular visual field of the observer at a fixed distance of 13 cm. An ascending and descending sequence of electric stimuli (0 µA, 50 µA, 150 µA, 300 µA, 500 µA, 0 µA, 500 µA, 300 µA, 150 µA, 50 µA; 1-second duration, with 60 seconds between stimuli) was delivered to the demonstrator. This sequence was repeated 6 times, resulting in 12 repetitions for each stimulation level. The total duration of the first part of the experiment was 1 hour.

To assess whether ECo operates through non-visual sensory mechanisms, an opaque visual occluder (black coated polystyrene) was placed between the two animals. After 5 minutes of habituation, electric stimuli (0 µA, 500 µA; 1-second duration, with 60 seconds between stimuli) were delivered to the demonstrator. The stimulation was repeated 12 times, resulting in 12 repetitions for each stimulation level. The total duration of the second part of the experiment was 20 minutes.

For the activity-dependent c-Fos staining experiment, mice were assigned to one of three groups: observers, demonstrators, and controls. Both the observer and demonstrator mice were head-fixed and positioned as described earlier, with stimulation electrodes attached. The experiment began with a 5-minute habituation period to let the mice acclimate to the context. Afterward, the demonstrators received an electric stimulus of 500 µA for 1 second, repeated every 60 seconds for a total of 5 shocks. The control mice were also head-fixed in the same setup for the same duration but did not receive or observe any shocks. Following the experiment, all mice were placed in isolated cages for 90 minutes to prevent any social interaction and allow for the peak expression of c-Fos (Morgan et al., 2005). After this 90-minutes period, the mice were transcardially perfused.

### Video recording

To further investigate whether ECo primarily relies on visual cues, we recorded videos of mice undergoing a simplified version of the aversive stimulation protocol used for demonstrators. These videos were then shown to observer mice. The recordings were made with an infrared USB camera connected to a Raspberry Pi 3, running a custom Python3 script with HEVC encoding in grayscale, and saved in .mp4 format at 20 fps. An infrared illuminator provided adequate lighting without altering the environmental luminance at visible wavelengths. Stimulation electrodes were attached to the tail of the mouse and connected to a stimulus isolator. The camera was positioned parallel to the mouse running direction, capturing the entire body from face to tail. To standardize the videos, the mouse eye was centered in a fixed area of the frame. Before the stimulation protocol began, a 20-minute video of baseline activity was recorded. The stimulation sequence consisted of an ascending and descending ladder of electric stimuli (0 µA, 50 µA, 150 µA, 300 µA, 500 µA, 0 µA, 500 µA, 300 µA, 150 µA, 50 µA; 1 second duration, with 60-second interstimulus intervals). This sequence was repeated three times, yielding a total of six repetitions per stimulation level.

### Video administration

Observer mice were head-fixed on a circular treadmill with a mock stimulating electrode at the base of their tail. They were placed 13 centimeters from the curved monitor used during habituation for head-fixation. The video of the demonstrator mouse was aligned so that its head was centered in the binocular visual field of the observer mouse. Mice underwent a 5-minute habituation period in which the observer watched a video of the unstimulated demonstrator. After the habituation, observer mice were shown randomized sequences of the same demonstrator. These sequences were presented in three post-processed conditions: the upright condition, where the video was shown as originally recorded; the inverted condition, where the video was flipped vertically; and the phase-scrambled condition, where the spatial structure of the video was disrupted by scrambling the Fourier phase, while preserving the physical properties. Videos of the demonstrator receiving 0 µA and 500 µA (20 sec pre-stimulation; 15 sec post-stimulation) tail shocks were administered to the observer, with each intensity randomly shown 18 times for every post-processed condition. The total duration of the experiment was 66 minutes. Pupillary diameter and locomotor activity of the observer were monitored for the entire duration of the experiment.

### Whole-brain c-Fos immunolabeling and tissue clearing

To evaluate whole-brain c-Fos expression mice were sacrificed 90 minutes after ECo experimental paradigm by transcardial perfusion with 0.09% saline solution followed by 4% paraformaldehyde (PFA) for tissue fixation. Brains were extracted, post-fixed for 3 hours in 4% PFA, washed three times in 1M Phosphate Buffer Solution (PBS) for one hour, and stored in PBS Na-Azide 0.02% solution at 4°C. iDISCO protocol (Renier et al., 2014, 2016) was used for whole-brain c-Fos immunolabeling and tissue clearing.

Samples were dehydrated in a methanol (MeOH) gradient: 20% MeOH, 40% MeOH, 60% MeOH, 80% MeOH, 100% MeOH, 100% MeOH in MilliQ (H_2_O) for 1h each at RT on a rotating wheel, before overnight delipidation in a solution of 33% MeOH / 66% dichloromethane (DCM, Sigma-Aldrich) at 4°C. Next, samples were rinsed twice in 100% MeOH for 1h, and bleached in 5% H_2_O_2_ in MeOH at 4°C overnight. Finally, samples were washed in 100% MeOH for 1h and gradually rehydrated in 80% MeOH, 60% MeOH, 40% MeOH, 20% MeOH (1h each, in MilliQ H_2_O) and PBS (twice for 1h). Samples were then incubated on an adjustable rotator at RT in a permeabilization solution of PBS containing 0.02% Na-Azide, 0.2% TritonX-100, 20% DMSO, 0.3M glycine for 2 days followed by incubation at RT in a blocking solution of PBS containing 0.02% Na-Azide, 0.2% TritonX-100, 10% DMSO, 6% Donkey Serum, for additional 2 days. Samples were then incubated in the primary antibody solution (Rabbit anti-c-Fos antibody, 1:5000, Cell Signalling, in PBS 0.02% Na-Azide, 0.2%, Tween, 10ug/mL Heparin, 5% DMSO and 3% Donkey Serum solution) on a rotating wheel at 37°C for 2 weeks. After primary antibody incubation brains were washed 6 times with a PBS solution containing 0.02% Na-Azide, 0.2% Tween, 10ug/mL Heparin. Next, samples were incubated with the secondary antibody (Donkey anti-Rabbit AlexaFluor 647, 1:800), in the same solution used for primary antibody, and placed at 37°C in rotation for 1 week. For tissue clearing, samples were dehydrated in a methanol gradient: 20% MeOH, 40% MeOH, 60% MeOH, 80% MeOH, 100% MeOH, 100% MeOH in MilliQ (H_2_O) for 1h each at RT on a rotating wheel and protected from the light. Delipidation was achieved with 3 hours incubation at RT in 33% MeOH / 66% DCM, followed by two washes of 20 minutes with 100% DCM. Samples were cleared in dibenzylether (DBE, Sigma-Aldrich) and protected from the light. Samples were stored in glass tubes in the dark at RT until imaging.

### Light-sheet fluorescence microscopy (LSFM)

3D whole-brain imaging was performed on the Ultramicroscope II (Miltenyi Biotec) equipped with a 4×/0.3NA objective and an Andor Zyla 5.5 sCMOS Camera. Samples were placed in an imaging chamber made of 100% quartz filled with DBE and illuminated from the side by the laser light. The light sheet was generated by a laser (wavelength 639 nm) and two cylindrical lenses, and the emitted signal was first filtered with a bandpass filter emission wavelength 680 nm. Laser power was set at 45% and the camera exposure time at 200.00 ms. ImspectorPro v7_1_4 software (Miltenyi Biotec) was used for image acquisition. The z-step between each image was fixed at 5 μm, and for tile imaging the overlap was set to 20%. Acquired volumes (16-bit tiff) had a radial resolution of 1.625 µm, and an axial resolution of 6.72 μm (NAL=L0.035). The resulting sequences of tiff files were processed with BigStitcher software (Hörl et al., 2019) to obtain a single stitched file. Stitched files were then imported into ClearMap2 toolbox (Kirst et al., 2020) to automate the detection of c-Fos^+^ cells.

### Automated c-Fos Cell detection and statistical analysis of area activation

Single stitched files (1 file /animal) were imported in ClearMap2 (Kirst et al., 2020) for atlas registration and cell detection. Images were first converted into npy format tridimensional matrix, (reporting pixel intensity on x and y and stacks on the z dimension) and then aligned to the Allen Brain Atlas. For atlas registration images were down-sampled to match raw data resolution (X: 1.625 μm, Y: 1.625 μm, Z: 5 μm) with the one of the reference atlas (X: 25 μm, Y: 25 μm, Z: 25 μm). Resampled data were aligned with the Allen Brain Atlas firstly through a linear (affine) transformation, and then optimized with a second non-linear (b-spline) transformation, using the Elastix software package (Klein et al., 2010; Shamonin et al., 2014). Next, cell detection was performed in four consecutive steps: 1) Background Removal; 2) Spot/Maxima Detection; 3) Cell Shape Detection (set to match the overall number of detected cells across animals); 4) Cell Size Detection (minimum pixel size selected: 7 pixel). Cell detection outputs include the number of detected cells, their spatial coordinates, and their fluorescence intensity. Since the registration of images to the Allen Mouse Brain Atlas can be executed across multiple hierarchical levels, the identification of activated cells can be performed within brain region, area, or subarea (including cortical layers). In this study, we selected 198 brain areas from the Allen Brain Atlas hierarchy (structure level= 8), excluding brain areas associated with the Pons, Medulla, Cerebellar cortex, and Cerebellar nuclei macro-areas. To identify brain areas differentially activated across conditions, we analyzed the density of detected cells (i.e., the raw count normalized by the brain area volume, as estimated by the Allen Brain Atlas). To compare two conditions, we employed the non-parametric Mann-Whitney U test (alpha=0.05).

Since values of c-Fos expression density provide a single activation index for each region per animal, functional connectivity estimation was performed by calculating covariance across subjects, rather than within subjects, across all pairs of brain areas. Next, functional connectivity matrices were created cross-correlating regional c-Fos expression densities across the Control, Demonstrator, and Observer groups. Statistically significant differences between conditions were assessed using the non-parametric Kruskal-Wallis test (alpha=0.05).

### c-Fos activation in Dyads

To assess the relationship between c-Fos activation in paired animals from the same dyad, we evaluated the linear correlation coefficient between paired animals in individual brain areas. We first paired animals based on the behavioral paradigm used to map brain activation, so that each observer was paired with the corresponding demonstrator used to induce emotional contagion. We then selected the 44 brain regions that showed a significant increase in c-Fos cell density in both the observer and demonstrator groups compared to the control. For each selected region, we evaluated the correlation coefficient between paired animals of the same dyad and compared the results with surrogate data generated by 42 different combinations of randomly shuffled paired animals across dyads. We set the threshold for significance at the 95th percentile of the distribution of shuffled data, and regions were considered to be significantly positively correlated only if they had a correlation coefficient greater than the threshold.

### Behavior and Pupillometry

Pupillary and locomotor time series were recorded at 20 Hz and manually inspected. Trials with saccades or large deflections caused by spontaneous changes in behavioral state were excluded from further analysis. In the pupillary trials, blinking activity was removed by applying a median filter to the absolute value of the first derivative of the pupil track, normalized from 0 to 1. Derivative values exceeding 0.2 were replaced with NaN and then linearly interpolated. The pupil track was smoothed using Savitzky-Golay filtering with a polynomial order of 6 and a window length of 1.5 seconds. Pupillary responses to each stimulation intensity were z-scored relative to a 1 seconds prestimulus window and baseline-corrected by subtracting the responses to the 0 µA shock condition. Locomotor activity was similarly smoothed using Savitzky-Golay filtering with the same polynomial order and window length. Responses were baseline-corrected using the average value from a 1-second prestimulus window, followed by subtracting the responses to the 0 µA shock condition. Peaks in both pupillary and locomotor responses were defined as the average value within a 1 to 2 seconds window after stimulus onset.

To evaluate the pupillary responses curves, we utilized a custom psychometric fitting method. This method involves fitting a sigmoid function to the data, to model the relationship between stimulus intensity and the pupillary response. The fitting process is performed with Python using nonlinear least squares optimization provided by the *scipy* library. The psychometric fit is executed in the following manner: the array of responses for each animal was averaged over the time period from stimulus onset (0 to 4 seconds). The arrays corresponding to the stimulus intensities were normalized, and the resulting pupillary responses were scaled between 0 and 1 for each animal. The fit is performed using the sigmoid function *C*(*C*) = *C +* (*C/*(*1 + C*(-*0*(*0-0_0_*)))) where *x* stimulus intensity, *x_0_* is the threshold, *k* is the slope of the curve, *L* is the upper asymptote, *b* is the baseline level of the curve. The threshold is defined as the stimulus intensity at which the pupillary response reaches its inflection point. The goodness was evaluated using the coefficient R^2^, which measures how well the fitted curve captures the variance in the data. To perform the ANOVA and repeated measures ANOVA we used the functions ‘anova’ and ‘rm_anova’, for post hoc tests we used the function ‘pairwise_tests’ with adjustment method Benjamini Hochberg FDR correction (Benjamini & Yekutieli, 2001) from Pingouin Python library (Vallat, 2018). Spearman correlations are performed using spearmanr from the Python package scipy (Virtanen et al., 2020).

### Anatomical Network analysis

Brain areas with significant activation with respect to control were used to construct the anatomical connectome network. The normalized connection density was obtained from a publicly available dataset of data driven high resolution mouse connectome (Knox et al., 2019). A weighted directed graph with all the activated areas was created using the NetworkX Python library. Leiden algorithm was calculated in iGraph (Csárdi et al., 2024) using the *leidenalg* Python library (https://github.com/vtraag/leidenalg) to find the partitions of the network based on the density of weighted directed connections (Traag et al., 2019). The Hyperlink-Induced Topic Search (HITS) algorithm was performed only in the areas activated in the observer. The HITS algorithm is primarily used for ranking web pages but can be applied to various networks to identify important nodes. The algorithm iteratively updates these scores to reflect the mutual reinforcement between hubs and authorities, providing insight into the network structure. Areas with high hub scores might represent regions that influence many other areas, acting as connectors or broadcasters of neural information. Conversely, areas with high authority scores may signify regions that are crucial recipients or integrators of information (Szczurek & Horeni, 2018). To evaluate the significance of hub and authority scores for specific brain areas, we implemented a bootstrap-based randomization procedure. Random networks were generated to create a null model for statistical comparison. At each iteration, a random network was generated by randomly selecting brain areas, preserving the total number of areas in the network. The target area of interest was included in every iteration, while the remaining nodes were randomly sampled from the set of brain areas analyzed using the Allen Brain Atlas. Connections from and to the same brain structure were removed. For each target brain area, 1,000 random networks were generated. In each iteration, hub and authority scores for the target area were calculated using the HITS algorithm. This process resulted in a distribution of random hub and authority scores for the target area, representing the expected scores under random conditions. We calculated the z-score to measure how many standard deviations the observed hub or authority score deviated from the mean of the bootstrap-generated distribution. The Z-score was then compared with the cumulative distribution function of the standard normal distribution to compute a one-tailed p-value.

## Data availability

All code for data analysis associated with the current paper is available at https://github.com/raffaelemazziotti/ECo_code

## Acknowledgements

We gratefully acknowledge NVIDIA Corporation’s support with the Jetson AGX Xavier Developer Kit donation for this research. This work was partially supported by AIRETT Associazione Italiana per la sindrome di Rett. Funded by the European Union -Next Generation EU Missione 4 Componente 2 Inv. 1.5 CUP I53C22000780001, and by Project “Tuscany Health Ecosystem – THE”, Spoke 8, granted by Next Generation EU – National Recovery and Resilience Plan (Piano Nazionale di Ripresa e Resilienza, NRRP) – Mission 4 Component 2 Investment 1.4 – Ministry of University and Research (MUR) Call N. 3277 Project Code ECS_00000017 MUR Directoral Decree n.1055, 23 June 2022, CUP B83C22003930001. PRIN2022 20228RMXBE to T.P. We would also like to thank Antonella Calvello, Vania Liverani and Renzo Di Renzo for technical assistance.

## Author Contribution

Conceptualization: RM, AV, MC, TP; Methodology: RM, MC, AV, TP, FC; Investigation: RM, AV, MC, TP, SZ, AS, SB, GS, RN, EMP; Visualization: RM, MC, SZ, AS, FC; Funding acquisition: TP, RM, SB ; Writing original draft: MC, SZ, AV, TP, SB, AS, RN, EMP, RM

## References

Adriaense, J. E. C., Martin, J. S., Schiestl, M., Lamm, C., & Bugnyar, T. (2019). Negative emotional contagion and cognitive bias in common ravens. Proceedings of the National Academy of Sciences of the United States of America, 116(23), 11547–11552.

Bekkali, S., Youssef, G. J., Donaldson, P. H., Albein-Urios, N., Hyde, C., & Enticott, P. G. (2021). Is the Putative Mirror Neuron System Associated with Empathy? A Systematic Review and Meta-Analysis. Neuropsychology Review, 31(1), 14–57.

Benjamini, Y., & Yekutieli, D. (2001). The control of the false discovery rate in multiple testing under dependency. Annals of Statistics, 29(4), 1165–1188.

Bible, E. (2013). Heightened emotional contagion in Alzheimer disease. Nature Reviews. Neurology, 9(7), 359–359.

Broly, P., & Deneubourg, J.-L. (2015). Behavioural Contagion Explains Group Cohesion in a Social Crustacean. PLoS Computational Biology, 11(6), e1004290.

Burbano Lombana, D. A., Macrì, S., & Porfiri, M. (2021). Collective Emotional Contagion in Zebrafish. Frontiers in Behavioral Neuroscience, 15, 730372.

Chang-Arana, Á. M., Surma-aho, A., Hölttä-Otto, K., & Sams, M. (2022). Under the umbrella: components of empathy in psychology and design. Design Science, 8, e20.

Choi, J., & Jeong, Y. (2017). Elevated emotional contagion in a mouse model of Alzheimer’s disease is associated with increased synchronization in the insula and amygdala. Scientific Reports, 7, 46262.

Csárdi, G., Nepusz, T., Müller, K., Horvát, S., Traag, V., Zanini, F., & Noom, D. (2024). igraph for R: R interface of the igraph library for graph theory and network analysis. Zenodo. 10.5281/ZENODO.7682609

Cuff, B. M. P., Brown, S. J., Taylor, L., & Howat, D. J. (2014). Empathy: A Review of the Concept. Emotion Review: Journal of the International Society for Research on Emotion. 10.1177/1754073914558466

Di Cesare, G., Di Dio, C., Marchi, M., & Rizzolatti, G. (2015). Expressing our internal states and understanding those of others. Proceedings of the National Academy of Sciences, 112(33), 10331–10335.

Ferencová, N., Višňovcová, Z., Bona Olexová, L., & Tonhajzerová, I. (2021). Eye pupil -a window into central autonomic regulation via emotional/cognitive processing. Physiological Research / Academia Scientiarum Bohemoslovaca, 70(Suppl4), S669–S682.

Flavell, S. W., Gogolla, N., Lovett-Barron, M., & Zelikowsky, M. (2022). The emergence and influence of internal states. Neuron, 110(16), 2545–2570.

Ganea, D. A., Bexter, A., Günther, M., Gardères, P.-M., Kampa, B. M., & Haiss, F. (2020). Pupillary Dilations of Mice Performing a Vibrotactile Discrimination Task Reflect Task Engagement and Response Confidence. Frontiers in Behavioral Neuroscience, 14, 159.

Giacomucci, G., Moschini, V., Piazzesi, D., Padiglioni, S., Caruso, C., Nuti, C., Munarin, A., Mazzeo, S., Galdo, G., Polito, C., Emiliani, F., Frigerio, D., Morinelli, C., Bagnoli, S., Ingannato, A., Nacmias, B., Sorbi, S., Berti, V., & Bessi, V. (2024). Disentangling empathy impairment along Alzheimer’s disease continuum: From subjective cognitive decline to Alzheimer’s dementia. Cortex; a Journal Devoted to the Study of the Nervous System and Behavior, 172, 125–140.

Grujic, N., Polania, R., & Burdakov, D. (2024). Neurobehavioral meaning of pupil size. Neuron. 10.1016/j.neuron.2024.05.029

Han, Y., Sichterman, B., Carrillo, M., Gazzola, V., & Keysers, C. (2020). Similar levels of emotional contagion in male and female rats. Scientific Reports, 10(1), 2763.

Harmsen, I. E. (2019). Empathy in Autism Spectrum Disorder. Journal of Autism and Developmental Disorders, 49(10), 3939–3955.

Helt, M. S., Fein, D. A., & Vargas, J. E. (2020). Emotional contagion in children with autism spectrum disorder varies with stimulus familiarity and task instructions. Development and Psychopathology, 32(1), 383–393.

Hernandez-Lallement, J., Gómez-Sotres, P., & Carrillo, M. (2022). Towards a unified theory of emotional contagion in rodents-A meta-analysis. Neuroscience and Biobehavioral Reviews, 132, 1229–1248.

Herrando, C., & Constantinides, E. (2021). Emotional Contagion: A Brief Overview and Future Directions. Frontiers in Psychology, 12, 712606.

Hörl, D., Rojas Rusak, F., Preusser, F., Tillberg, P., Randel, N., Chhetri, R. K., Cardona, A., Keller, P. J., Harz, H., Leonhardt, H., Treier, M., & Preibisch, S. (2019). BigStitcher: reconstructing high-resolution image datasets of cleared and expanded samples. Nature Methods, 16(9), 870–874.

Jabarin, R., Levy, N., Abergel, Y., Berman, J. H., Zag, A., Netser, S., Levy, A. P., & Wagner, S. (2021). Pharmacological modulation of AMPA receptors rescues specific impairments in social behavior associated with the A350V Iqsec2 mutation. Translational Psychiatry, 11(1), 1–11.

Kareklas, K., & Oliveira, R. F. (2024). Emotional contagion and prosocial behaviour in fish: An evolutionary and mechanistic approach. Neuroscience and Biobehavioral Reviews, 163, 105780.

Keysers, C., & Gazzola, V. (2023). Vicarious Emotions of Fear and Pain in Rodents. Affective Science, 4(4), 662–671.

Keysers, C., Knapska, E., Moita, M. A., & Gazzola, V. (2022). Emotional contagion and prosocial behavior in rodents. Trends in Cognitive Sciences, 26(8), 688–706.

Kimmig, A.-C. S., Burger, L., Schall, M., Derntl, B., & Wildgruber, D. (2024). Impairment of affective and cognitive empathy in high functioning autism is mediated by alterations in emotional reactivity. Scientific Reports, 14(1), 21662.

Kim, S.-W., Kim, M., Baek, J., Latchoumane, C.-F., Gangadharan, G., Yoon, Y., Hyung, K. D.-S. L., & Shin, H.-S. (2023). Hemispherically lateralized rhythmic oscillations in the cingulate-amygdala circuit drive affective empathy in mice. Neuron, 111(3), 418–429.e4.

Kim, S.-W., Kim, M., & Shin, H.-S. (2021). Affective empathy and prosocial behavior in rodents. Current Opinion in Neurobiology, 68, 181–189.

Kirst, C., Skriabine, S., Vieites-Prado, A., Topilko, T., Bertin, P., Gerschenfeld, G., Verny, F., Topilko, P., Michalski, N., Tessier-Lavigne, M., & Renier, N. (2020). Mapping the fine-scale organization and plasticity of the brain vasculature. Cell, 180(4), 780–795.e25.

Klasen, M., Chen, Y.-H., & Mathiak, K. (2012). Multisensory emotions: perception, combination and underlying neural processes. Reviews in the Neurosciences, 23(4), 381–392.

Klein, A. S., Dolensek, N., Weiand, C., & Gogolla, N. (2021). Fear balance is maintained by bodily feedback to the insular cortex in mice. Science, 374(6570), 1010–1015.

Kleinberg, J. M. (1999). Authoritative sources in a hyperlinked environment. Journal of the ACM, 46(5), 604–632.

Knox, J. E., Harris, K. D., Graddis, N., Whitesell, J. D., Zeng, H., Harris, J. A., Shea-Brown, E., & Mihalas, S. (2019). High-resolution data-driven model of the mouse connectome. *Network Neuroscience (Cambridge*, Mass.), 3(1), 217–236.

Kutlubaev, M. A., Shagieva, D. R., Karimova, G. I., Izmalkova, A. I., & Myachikov, A. V. (2024). Pupillometry in the Assessment of Psychoemotional State and Cognitive Functions in Humans. Neuroscience and Behavioral Physiology, 54(1), 112–121.

Laeng, B., Sirois, S., & Gredebäck, G. (2012). Pupillometry: A Window to the Preconscious? Perspectives on Psychological Science: A Journal of the Association for Psychological Science, 7(1), 18–27.

Lamm, C., Bukowski, H., & Silani, G. (2016). From shared to distinct self–other representations in empathy: evidence from neurotypical function and socio-cognitive disorders. Philosophical Transactions of the Royal Society of London. Series B, Biological Sciences. 10.1098/rstb.2015.0083

Mazziotti, R., Carrara, F., Viglione, A., Lupori, L., Lo Verde, L., Benedetto, A., Ricci, G., Sagona, G., Amato, G., & Pizzorusso, T. (2021). MEYE: Web App for Translational and Real-Time Pupillometry. eNeuro, 8(5). 10.1523/ENEURO.0122-21.2021

McGinley, M. J., David, S. V., & McCormick, D. A. (2015). Cortical Membrane Potential Signature of Optimal States for Sensory Signal Detection. Neuron, 87(1), 179–192.

Meyza, K., & Knapska, E. (2018). What can rodents teach us about empathy? Current Opinion in Psychology, 24, 15–20.

Mondoloni, S., Molina, P., Lecca, S., Wu, C.-H., Michel, L., Osypenko, D., Cachin, F., Flanigan, M., Congiu, M., Lalive, A. L., Kash, T., Deng, F., Li, Y., & Mameli, M. (2024). Serotonin release in the habenula during emotional contagion promotes resilience. Science. 10.1126/science.adp3897

Morgan, J. I., Cohen, D. R., Hempstead, J. L., & Curran, T. (2005). Mapping behaviorally relevant neural circuits with immediate-early gene expression. Current Opinion in Neurobiology, 15(5), 599–606.

Nakakoga, S., Higashi, H., Muramatsu, J., Nakauchi, S., & Minami, T. (2020). Asymmetrical characteristics of emotional responses to pictures and sounds: Evidence from pupillometry. PloS One, 15(4), e0230775.

Nicolas, C., Ju, A., Wu, Y., Eldirdiri, H., Delcasso, S., Couderc, Y., Fornari, C., Mitra, A., Supiot, L., Vérité, A., Masson, M., Rodriguez-Rozada, S., Jacky, D., Wiegert, J. S., & Beyeler, A. (2023). Linking emotional valence and anxiety in a mouse insula-amygdala circuit. Nature Communications, 14(1), 1–18.

Pallett, P. M., & Meng, M. (2015). Inversion effects reveal dissociations in facial expression of emotion, gender, and object processing. Frontiers in Psychology, 6, 1029.

Pisansky, M. T., Hanson, L. R., Gottesman, I. I., & Gewirtz, J. C. (2017). Oxytocin enhances observational fear in mice. Nature Communications, 8(1), 1–11.

Plata-Bello, J., Privato, N., Modroño, C., Pérez-Martín, Y., Borges, Á., & González-Mora, J. L. (2023). Empathy Modulates the Activity of the Sensorimotor Mirror Neuron System during Pain Observation. Behavioral Sciences, 13(11), 947.

Poltoratski, S., Kay, K., Finzi, D., & Grill-Spector, K. (2021). Holistic face recognition is an emergent phenomenon of spatial processing in face-selective regions. Nature Communications, 12(1), 4745.

Prochazkova, E., & Kret, M. E. (2017). Connecting minds and sharing emotions through mimicry: A neurocognitive model of emotional contagion. Neuroscience and Biobehavioral Reviews, 80, 99–114.

Qi, G., Zhang, P., Li, T., Li, M., Zhang, Q., He, F., Zhang, L., Cai, H., Lv, X., Qiao, H., Chen, X., Ming, J., & Tian, B. (2022). NAc-VTA circuit underlies emotional stress-induced anxiety-like behavior in the three-chamber vicarious social defeat stress mouse model. Nature Communications, 13(1), 1–19.

Renier, N., Adams, E. L., Kirst, C., Wu, Z., Azevedo, R., Kohl, J., Autry, A. E., Kadiri, L., Umadevi Venkataraju, K., Zhou, Y., Wang, V. X., Tang, C. Y., Olsen, O., Dulac, C., Osten, P., & Tessier-Lavigne, M. (2016). Mapping of Brain Activity by Automated Volume Analysis of Immediate Early Genes. Cell, 165(7), 1789–1802.

Renier, N., Wu, Z., Simon, D. J., Yang, J., Ariel, P., & Tessier-Lavigne, M. (2014). iDISCO: a simple, rapid method to immunolabel large tissue samples for volume imaging. Cell, 159(4), 896–910.

Rizzolatti, G., Fadiga, L., Gallese, V., & Fogassi, L. (1996). Premotor cortex and the recognition of motor actions. Brain Research. Cognitive Brain Research, 3(2), 131–141.

Scheggia, D., Managò, F., Maltese, F., Bruni, S., Nigro, M., Dautan, D., Latuske, P., Contarini, G., Gomez-Gonzalo, M., Requie, L. M., Ferretti, V., Castellani, G., Mauro, D., Bonavia, A., Carmignoto, G., Yizhar, O., & Papaleo, F. (2019). Somatostatin interneurons in the prefrontal cortex control affective state discrimination in mice. Nature Neuroscience, 23(1), 47–60.

Silasi, G., Xiao, D., Vanni, M. P., Chen, A. C. N., & Murphy, T. H. (2016). Intact skull chronic windows for mesoscopic wide-field imaging in awake mice. Journal of Neuroscience Methods, 267, 141–149.

Smith, M. L., Asada, N., & Malenka, R. C. (2021). Anterior cingulate inputs to nucleus accumbens control the social transfer of pain and analgesia. Science. 10.1126/science.abe3040

Solié, C., Contestabile, A., Espinosa, P., Musardo, S., Bariselli, S., Huber, C., Carleton, A., & Bellone, C. (2022). Superior Colliculus to VTA pathway controls orienting response and influences social interaction in mice. Nature Communications, 13(1), 1–15.

Szczurek, P., & Horeni, M. (2018). Using link analysis algorithms to study the role of neurons in the worm connectome. 2018 IEEE 32nd International Conference on Advanced Information Networking and Applications (AINA), 5, 651–657.

Traag, V. A., Waltman, L., & van Eck, N. J. (2019). From Louvain to Leiden: guaranteeing well-connected communities. Scientific Reports, 9(1), 5233.

Vallat, R. (2018). Pingouin: statistics in Python. Journal of Open Source Software, 3(31), 1026.

Viglione, A., Mazziotti, R., & Pizzorusso, T. (2023). From pupil to the brain: New insights for studying cortical plasticity through pupillometry. Frontiers in Neural Circuits, 17, 1151847.

Vinck, M., Batista-Brito, R., Knoblich, U., & Cardin, J. A. (2015). Arousal and locomotion make distinct contributions to cortical activity patterns and visual encoding. Neuron, 86(3), 740– 754.

Virtanen, P., Gommers, R., Oliphant, T. E., Haberland, M., Reddy, T., Cournapeau, D., Burovski, E., Peterson, P., Weckesser, W., Bright, J., van der Walt, S. J., Brett, M., Wilson, J., Millman, K. J., Mayorov, N., Nelson, A. R. J., Jones, E., Kern, R., Larson, E., … SciPy 1.0 Contributors. (2020). SciPy 1.0: fundamental algorithms for scientific computing in Python. Nature Methods, 17(3), 261–272.

Wang, D., Liu, C., & Chen, W. (2024). The role of self-representation in emotional contagion. Frontiers in Human Neuroscience, 18, 1361368.

Warren, A. (2022). Heightened emotion processing as a compensatory mechanism in persons with Alzheimer’s disease: Psychological insights from the tri-network model. Frontiers in Dementia, 1, 983331.

Wenig, K., Boucherie, P. H., & Bugnyar, T. (2021). Early evidence for emotional play contagion in juvenile ravens. Animal Cognition, 24(4), 717–729.

Wu, W.-Y., Cheng, Y., Liang, K.-C., Lee, R. X., & Yen, C.-T. (2023). Affective mirror and anti-mirror neurons relate to prosocial help in rats. iScience, 26(1), 105865.

Yu, D., Bao, L., & Yin, B. (2024). Emotional contagion in rodents: A comprehensive exploration of mechanisms and multimodal perspectives. Behavioural Processes, 216, 105008.

Zheng, C., Huang, Y., Bo, B., Wei, L., Liang, Z., & Wang, Z. (2020). Projection from the Anterior Cingulate Cortex to the Lateral Part of Mediodorsal Thalamus Modulates Vicarious Freezing Behavior. Neuroscience Bulletin, 36(3), 217–229.

