## Supplementary figures and images for "Pupillometry and Whole-Brain c-Fos Mapping Uncover Multimodal Mirror Mechanisms in Emotional Contagion Networks of Mice"

### S1.png

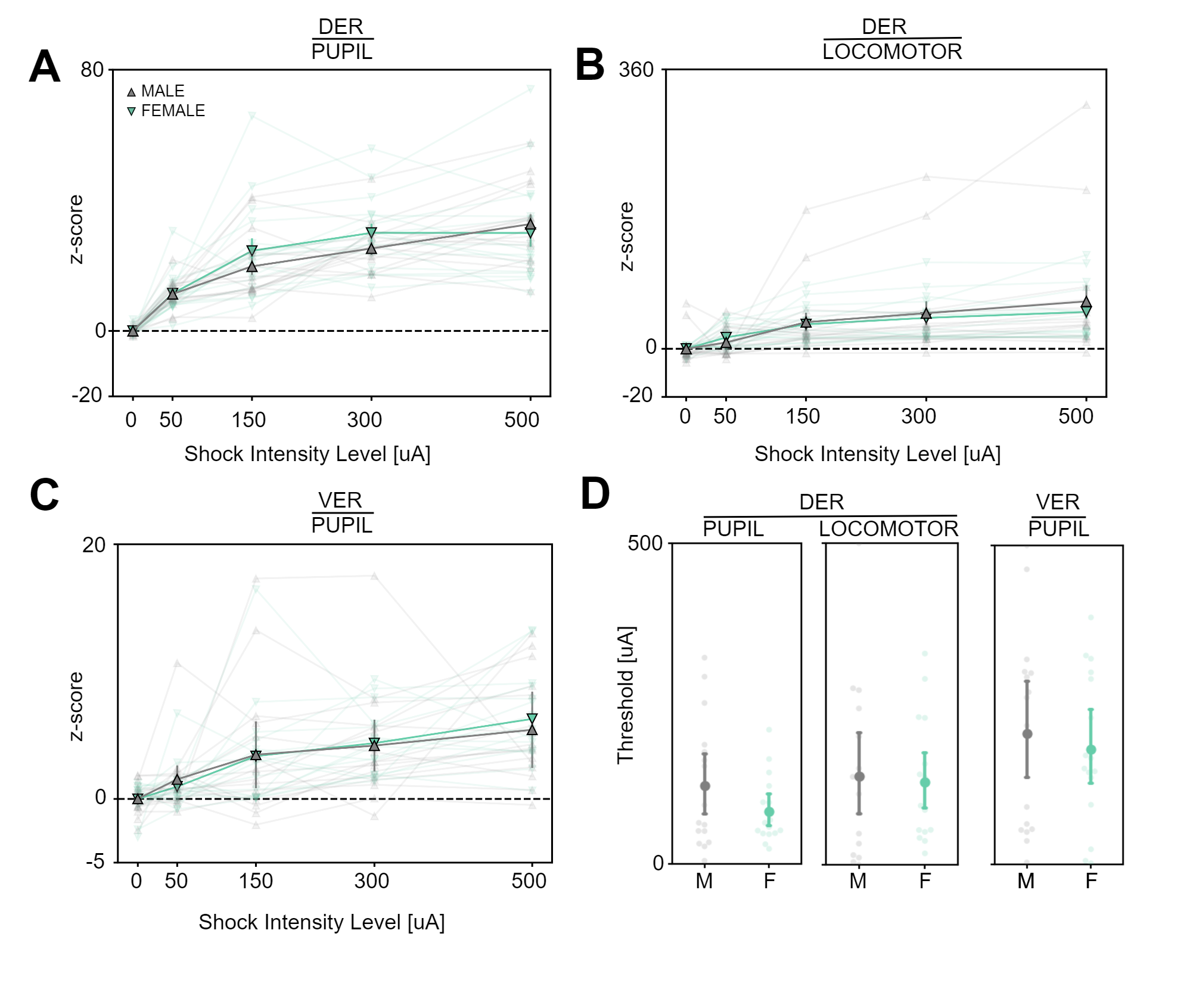

### S2.png

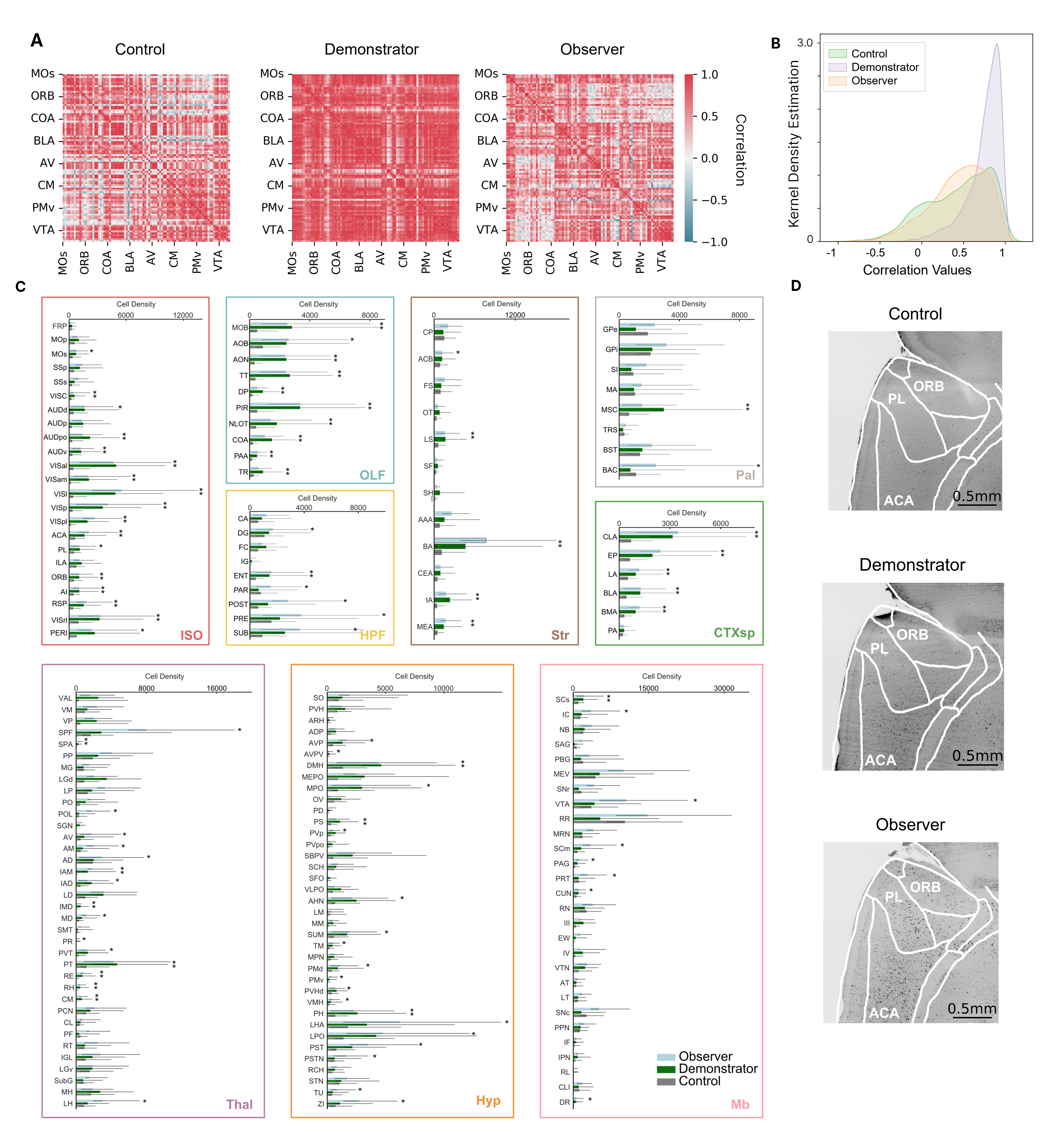
